# COVID-19 Contact Tracing Analysis with Bluetooth Technology Using Raspberry Pis

**DOI:** 10.1101/2022.08.23.504944

**Authors:** Dean Zhang

## Abstract

Contact tracing, a method for detecting and preventing the spread of a disease, can become more efficient by becoming automated, rather than being done manually. Bluetooth based contact tracing is a potential method for creating an automated method for contact tracing. However, Bluetooth signals cannot always predict something with complete accuracy due to many subtle obstructions. Received signal strength indicator (RSSI), a value produced when Bluetooth devices send and receive signals, generated from Raspberry Pis can predict the distance between the two devices. By running many experiments with variations of obstructions, I was able to successfully create models to correlate RSSI values and distance with a potential success rate of 91.973%. For my research, a success is determined to be anything but a false negative. Despite the limitations when conducting my research, the inaccuracies of my results prove Bluetooth based contact tracing to not be a reliable method for determining people who have been in contact with an index case in the real world.

## I. Introduction

### A. Project Description

My project focuses on using Bluetooth signals from Raspberry Pis to predict the distance between the two devices when different obstructions interfere. As there are countless obstructions everywhere, it is crucial to determine if distance can be predicted from Bluetooth signals. Ultimately, with that ability and the ability to predict a duration of time, this project relates to piPACT’s goal of contact tracing. Moreover, anonymity is also a crucial part to piPACT’s goals; therefore, using the different forms of identification generated by the raspberry pis, I can operate under a private setting. In general, I used common items that may block a signal from leaving a device one carries like shorts, jeans, a human body, and more to. From there, I simply made a model and a basic user interface to test the accuracy and privacy of my work.

### B. Background Information

The devices I used are Raspberry Pis. For the purpose of this project, they are simply devices that advertise and scan Bluetooth signals [1]. Bluetooth signals are signals that operate at about a 2.4 gigahertz frequency to transmit information from once device to another [2]. Specifically, the received signal strength indicator (RSSI), a value to convey the strength of the received Bluetooth signal, is a value generated from the Bluetooth signals. Both physical and atmospheric changes in the setting affect the RSSI value [3]. From RSSI, it is possible to predict distance from RSSI, which can then be used for contact tracing [4]. Contact tracing is a method to identify and contact the people who have been too close to someone with a disease for too long [5]. For the current COVID-19 pandemic, any distance less than or equal to six feet is classified to be too close, and any period of time greater or equal to ten minutes is considered too long [6]. For my research, I assumed that Bluetooth signals will solely be transmitted from Raspberry Pis in an indoor setting that is extremely similar to my room. Although uncontrollable, I do recognize the atmospheric changes that occurred when I was collecting data. Moreover, many more subtle things like sound or Wi-Fi interferences may have affected the Bluetooth signals. Adding on, as Bluetooth signals may bounce off of objects, the objects in the setting that were in close proximity to the Raspberry Pis could also have altered the signal [2].

## II. Hypothesis/Hypotheses

Hypothesis 1 is physical obstructions will decrease the RSSI values, thus affecting proximity prediction. Specifically, I am interested to see how much a given type of obstruction affects the RSSI value, so a better prediction of distance can be made. Hypothesis 2 is atmospheric obstructions will decrease the RSSI values, thus allowing for a better prediction of distance. Hypothesis 3 is that it is possible to create a detection algorithm from Bluetooth signals generated between two Raspberry Pis. The ultimate goal would be to not just create an algorithm, but to create an accurate one. Hypothesis 4 is putting the same obstruction on the scanner Raspberry Pi will result in an insignificant difference of RSSI values when compared to the RSSI values generated from putting the same obstruction on the advertising Raspberry Pi.

- All hypotheses result in the ability to better predict distance and duration of time at which the two devices coexist at, which can later be used for contact tracing.
- Because the measured variable is a Bluetooth signal value and the predicted variables, distance and time, are crucial for contact tracing, the hypotheses are directly related to Bluetooth based contact tracing.
- For hypothesis 1, the specific obstructions observed are a pair of athletic shorts, a pair of jeans, a cabinet/shelf, and a human body.
- Hypothesis 2 tests the effects of temperature (Fahrenheit) and humidity (relative humidity).
- Hypothesis 3 focuses on creating a model and algorithm that accurately links RSSI values to distance for future distance prediction.
- Hypothesis 4 tests if a pair of shorts or jeans blocking the advertising Raspberry Pi results in approximately equally RSSI values as to when the same pair of shorts or jeans is blocking the scanner Raspberry Pi.

## III. Experiments and Data Collections

The only variables I will be directly changing is distance and type of obstruction, meaning everything else that is in my control will remain constant. To be a little more specific about a variation, for a pair of shorts, I will conduct one experiment by placing a pair of shorts of one Raspberry pi, the on another Raspberry Pi, then on both Raspberry Pis. Additionally, every time I advertise Bluetooth signals from the advertising Raspberry Pi, I will take note of the temperature and humidity at that moment too.

### A. Plan and Execution

I collected data by running my Raspberry Pis for three minutes at a distance while recording the temperature and humidity. Then, I would increase the distance by 3.5 inches from 0 inches to 143.5 inches. Additionally, I did measure the RSSI values when the Raspberry Pis were at 72 and 144 inches. To ensure reproducibility, for all experiments, the Raspberry Pis were orientated the same way. They were placed in the same room, so the amount of signal bouncing off surfaces each experiment should be approximately equal. They were on the same surface, the floor, except when they were in pockets of clothing. When there were in pockets, I made sure to fully surround the device so ensure a minimal amount of singal escaping through a path that is obstruction-free. The advertiser Raspberry Pi’s TX power was consistent throughout. The varied distances were pre-measured, so there should be little to no variation within the changing of distance for the experiments.

There were changes in weather, and I tried to take that into account by recording the temperature and humidity every time I ran an experiment. The interference of signals coming from other devices, like Wi-Fi signals or sound were not well controlled during the experiments. Overall, simply running the experiments in a room full of objects, which can lead to signals bouncing off object, was a limitation I couldn’t overcome. Despite clearing a direct path between the two Raspberry Pis, I was unable to provide a clean and object-free environment for my experimentation.

### B. Data Relevance

The goal of this research project for me was to simulate real life situations when predicting distance from Bluetooth signals. Therefore, the physical obstructions in each experiment were chosen because they commonly applied in the outside world: clothes as Bluetooth devices are in pockets, a shelf as grocery shelves or bookshelves are common, and humans as phones could be in a back pocket or a backpack. Therefore, these obstructions were perfect for testing to see how those obstructions affect RSSI values, as they are commonly used in the real world. Additionally, atmospheric values were measured to test hyp. 2. Ultimately, the data collected is going to be used to create an algorithm that can predict distance and time to a certain degree of accuracy. Experiments 2, 3, 5, and 6 were conducted to see whether the Raspberry Pi that experienced the obstruction mattered.

### C. Examples

## IV. V. Analysis and Algorithms

### A. Description

First off, I performed a paired t test on the exp. 2&3 and 5&6 to test hyp. 4. Specifically, I subtracted the data of exp. 3 and exp. 6 from exp. 2 and exp. 5, respectively. Likewise, I performed a 2–sample t test on all experiments and exp. 1 to test hyp. 1. Additionally, to determine whether putting an obstruction on one Raspberry Pi is the same as putting that obstruction on both Raspberry Pis, I did a 2-sample t-test on experiments 2 & 4, 3 & 4, 5 & 7, and 6 & 7. This was to test if there even is a difference between those two two scenarios.

Secondly, the goal of my algorithms was to ultimately be able to predict the distance given an RSSI value, the temperature, the humidity, and the type of obstruction. Because I took upon three independent variables (distance, temperature, and humidity) at once, I created a multiple regression, a multivariable linear regression. In order to make the data linear, I first took the natural log of all the x values (distance, temperature, and humidity). In order to avoid skewing the calculations too much, I removed all data at distance 0.0 inches due to the problems resulting from taking the natural log of 0. From there on, a regression was made to ultimately predict distance from the other three variables. Similarly, I made another regression but without temperature and humidity to test hyp. 2, to see if atmospheric changes really affect RSSI values in a way that make predicting distance more accurate.

Thirdly, to compare the results, I ran my data through the models I made to determine the number of true & false positives and true & false negatives I would get in a real-life scenario.

Lastly, based on the universally unique identifies (UUID), major and minor values generated by the Raspberry Pis, I created some functions to compare and see whether or not those values from a data file math a fake index case. Similarly, based on the timestamps on the file names, I was able to successfully measure the amount of time spent in between two devices. Then, I created a simple user interface to run not just my distance predictions, but also to test the anonymity and period of time spent in contact.

Overall, I chose to use Python due to its abundance of capabilities and resources when it comes to dealing with regressions and plots. Also, because the code written to control the Raspberry Pis is in Python, I thought having my code in Python would be helpful to serve as conformity.

### B. Results and Examples

First off, it is apparent that in table 2, many statistical tests resulted in a large p-value compared to alpha, meaning a statistically insignificant difference between two datasets. Therefore, for all tested obstructions, except for exp. 9, they could all belong to the same population of data. Also, putting an obstruction on one Raspberry Pi led to no significant difference compared to putting that obstruction on both Raspberry Pis. For example, exp. 3 & 4 resulted in high p-values that made the statistically insignificant to put shorts on one Raspberry Pi and on both Raspberry Pis. The same goes for jeans.

**TABLE I.**
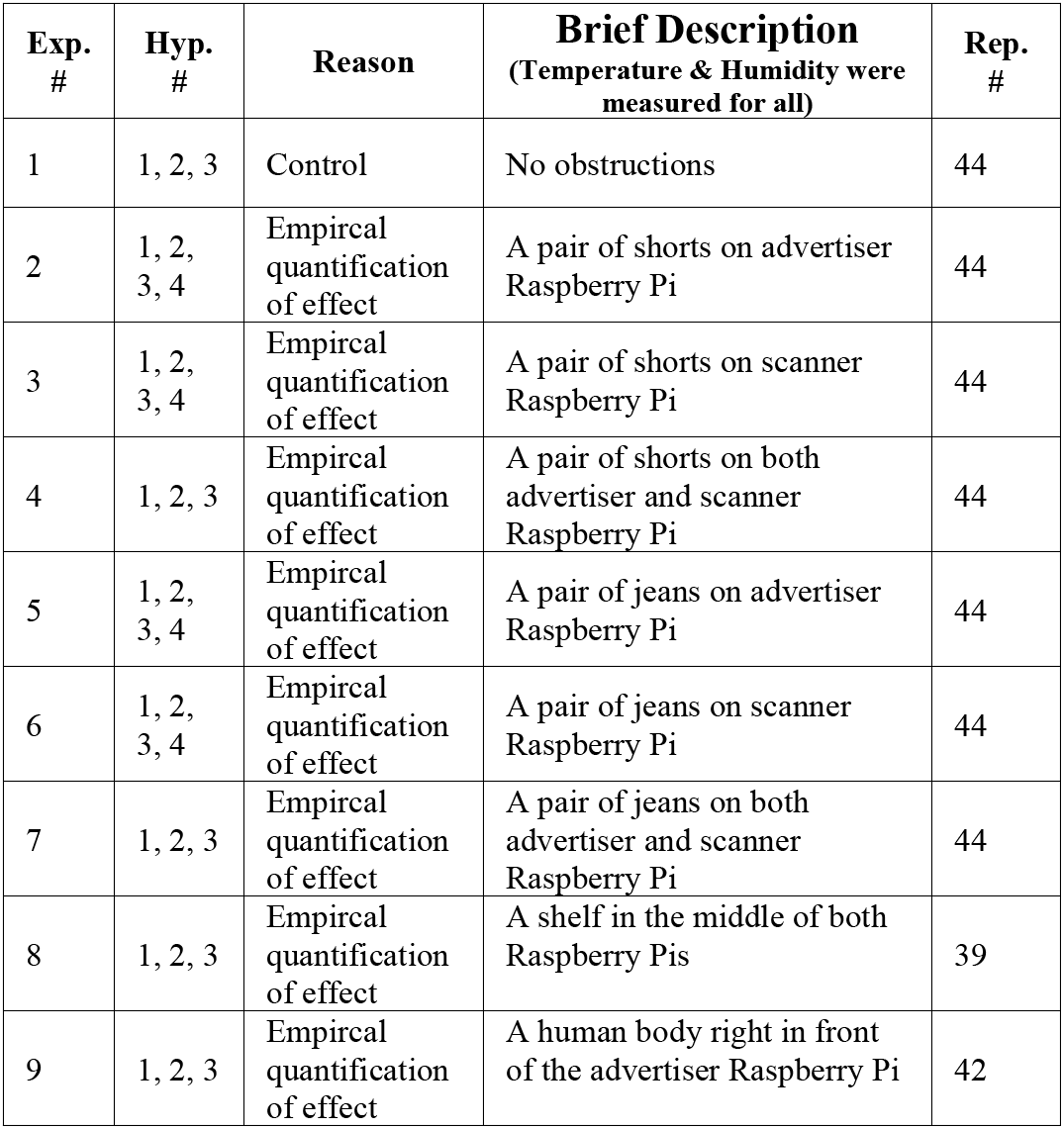
EXPERIMENTS OVERVIEW.

| Exp. # | Hyp. # | Reason | Brief Description<br>(Temperature & Humidity were measured for all) | Rep. # |
| --- | --- | --- | --- | --- |
| 1 | 1, 2, 3 | Control | No obstructions | 44 |
| 2 | 1, 2, 3, 4 | Empirical quantification of effect | A pair of shorts on advertiser Raspberry Pi | 44 |
| 3 | 1, 2, 3, 4 | Empirical quantification of effect | A pair of shorts on scanner Raspberry Pi | 44 |
| 4 | 1, 2, 3 | Empirical quantification of effect | A pair of shorts on both advertiser and scanner Raspberry Pi | 44 |
| 5 | 1, 2, 3, 4 | Empirical quantification of effect | A pair of jeans on advertiser Raspberry Pi | 44 |
| 6 | 1, 2, 3, 4 | Empirical quantification of effect | A pair of jeans on scanner Raspberry Pi | 44 |
| 7 | 1, 2, 3 | Empirical quantification of effect | A pair of jeans on both advertiser and scanner Raspberry Pi | 44 |
| 8 | 1, 2, 3 | Empirical quantification of effect | A shelf in the middle of both Raspberry Pis | 39 |
| 9 | 1, 2, 3 | Empirical quantification of effect | A human body right in front of the advertiser Raspberry Pi | 42 |

**TABLE II.**
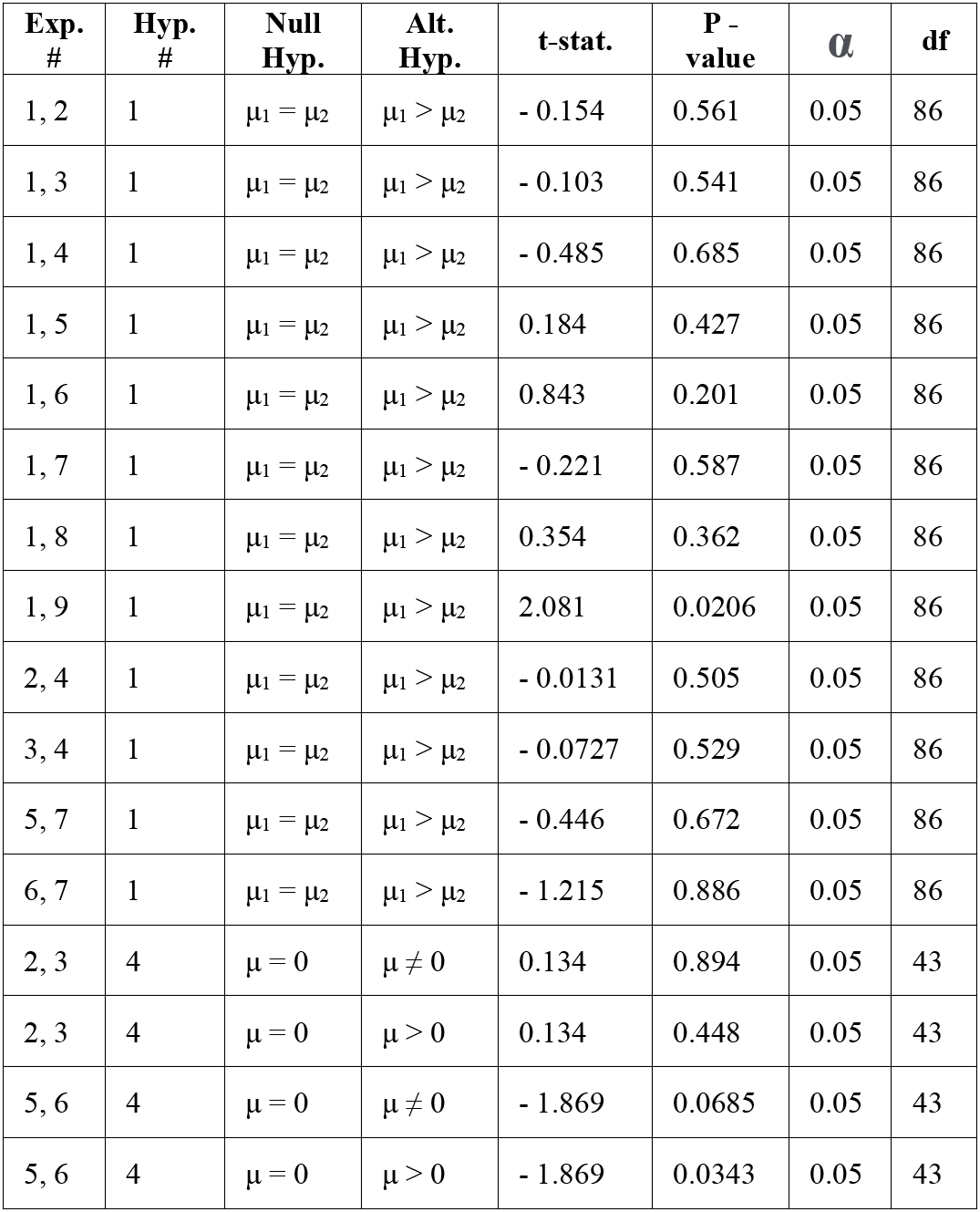
STATISTICAL SIGNIFICANT TESTS ON EXPERIMENTS.

Before transforming the data into a linear form, I plotted what the relationship between RSSI values and distance is supposed to look like. Figure 3 makes it apparent that the relationship between the two are a type of logarithmic function as the curve is consistent, no matter the experiment is being done.

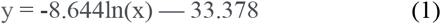

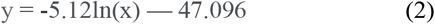

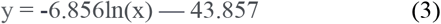

In equations 1, 2, and 3, y is the measured RSSI value, and x represents distance. Equation one is based on exp 1., (2) is based on exp. 4, and (3) is based on exp. 9. Equation 1, 2, and 3 are examples of the true regression model between RSSI and distance, when nothing is transformed. Just Equation 1 has highest intercept and (2) has the smallest slope (when comparing the absolute values of all slopes), suggesting that it has the least intense drop at the beginning of its logarithmic curve. Due to the logarithmic nature of a curve, it could be inferred that a sudden and dramatic change in RSSI values, could suggest that the distance is well below the six-feet threshold, assuming the change is not caused by an obstruction. Additionally, out of all the experiments, only exp. 9 was proven to statistically significant when compared to exp. 1. It apparent as the curve for exp. 9 is consistently below all the other curves, suggesting that it has had the most effect on obstructing the Bluetooth signal.

After building the model, I was able to test it. I ended up getting the number of positives and negatives my model would get if it were diagnosing people. Because all except exp. 9 were statistically insignificantly different from exp. 1, I tested the from one experiment with models generated from another experiment. So, to further support the insignificance, it is clear in table 3 that there are cases where a model from an experiment can be applied on another experiment and lead to better results compared to when that experiment’s model was applied on itself. For example, exp. 8 resulted in the same four numbers when put through the model made from exp. 4 and exp. 7. The same goes with how exp. 9 put through the model created from exp. 4 and exp. 7 resulted in the same four numbers.

**TABLE III.** PROXIMITY PREDICTIONS SOLEY BASED ON RSSI VALUES.

| Exp. # | Model from Exp. # | True Pos. | False Pos. | True Neg. | False Neg. |
| --- | --- | --- | --- | --- | --- |
| 1 | 1 | 16 | 7 | 15 | 6 |
| 4 | 4 | 15 | 7 | 16 | 6 |
| 4 | 8 | 22 | 14 | 8 | 0 |
| 7 | 7 | 16 | 9 | 13 | 6 |
| 7 | 8 | 19 | 13 | 9 | 3 |
| 7 | 9 | 21 | 16 | 6 | 1 |
| 8 | 4 | 8 | 1 | 22 | 8 |
| 8 | 7 | 8 | 1 | 22 | 8 |
| 8 | 8 | 12 | 6 | 16 | 5 |
| 9 | 4 | 6 | 0 | 23 | 13 |
| 9 | 7 | 6 | 0 | 23 | 13 |
| 9 | 9 | 15 | 4 | 18 | 5 |

Sometimes another experiment’s model led to even better results. In this case, better results mean that there are more true positive and true negatives. However, as false negatives are considered worse than false positives in my project, a significant number of false negatives automatically is considered bad results. For example, exp. 4 put through model from exp. 8 led to zero false negatives. Although, the number of true positives increased, I consider that a success. A true positive at most means a more severe case of self-isolating. Solely, based off this one situation, it can be interpreted that no one should spread the disease further because there are no false negatives. The same situation is present when exp. 7 is passed through the model made my exp. 9.

iMow, ι uaiisiνimcα me lνgaiuiuiuu uuivc miν a uiicai one and took into account temperature and humidity. Similar to how exp. 9’s curve is consistently below all the other curves, exp. 9’s linear line is also below all the other lines, suggesting that exp. 9 is truly significantly different from the rest.

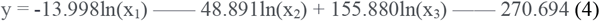

In (4), y = a measured RSSI value, x_1_ is distance, x_2_ is temperature, and x3 is humidity. Equation 4 is not plotted, but that is an example of the equation I used when calculating the numbers in table 4. Figure 4 is simply just a graph of the transformed data originally presented in figure 3. As all the coefficients are the same, the same trends in slope and intercept are visible.

**TABLE IV.** PROXIMITY PREDICTIONS BASED ON RSSI, TEMPERATURE, AND HUMIDITY.

| Exp. # | Model from Exp. # | True Pos. | False Pos. | True Neg. | False Neg. |
| --- | --- | --- | --- | --- | --- |
| 1 | 1 | 19 | 3 | 20 | 2 |
| 4 | 4 | 5 | 0 | 23 | 16 |
| 4 | 8 | 13 | 6 | 17 | 8 |
| 7 | 1 | 5 | 0 | 22 | 17 |
| 7 | 4 | 5 | 0 | 22 | 17 |
| 7 | 7 | 9 | 0 | 23 | 12 |
| 7 | 8 | 10 | 6 | 16 | 12 |
| 8 | 8 | 13 | 7 | 15 | 4 |
| 9 | 7 | 6 | 0 | 23 | 13 |
| 9 | 8 | 6 | 0 | 23 | 13 |
| 9 | 9 | 14 | 8 | 15 | 5 |

**Fig. 1.**
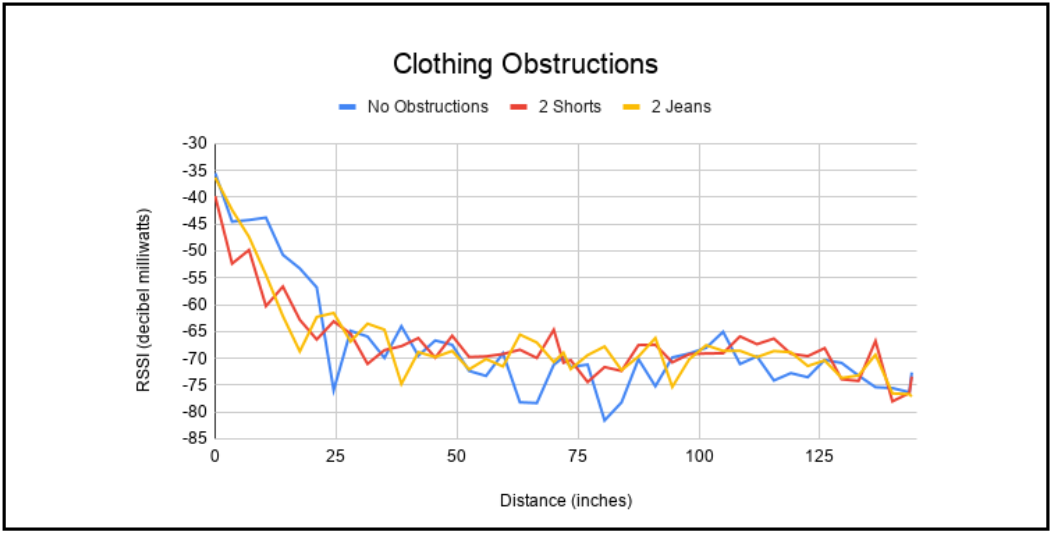
The comparison of the relationship between RSSI and distance for exp. 1, 4, and 7.

**Fig. 2.**
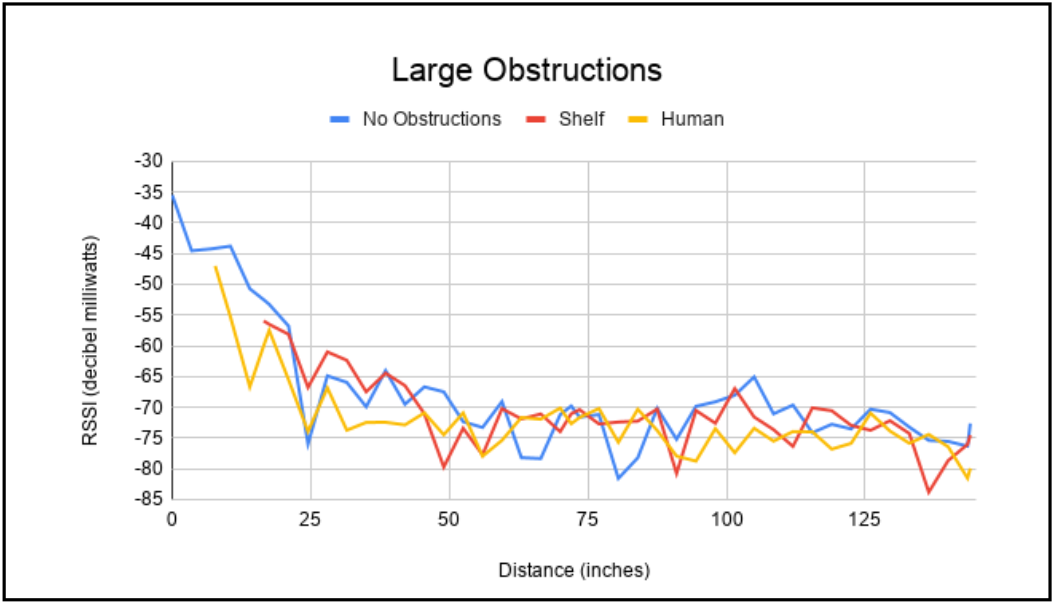
The comparison of the relationship between RSSI and distance for exp. 1, 8, and 9.

**Fig. 3.**
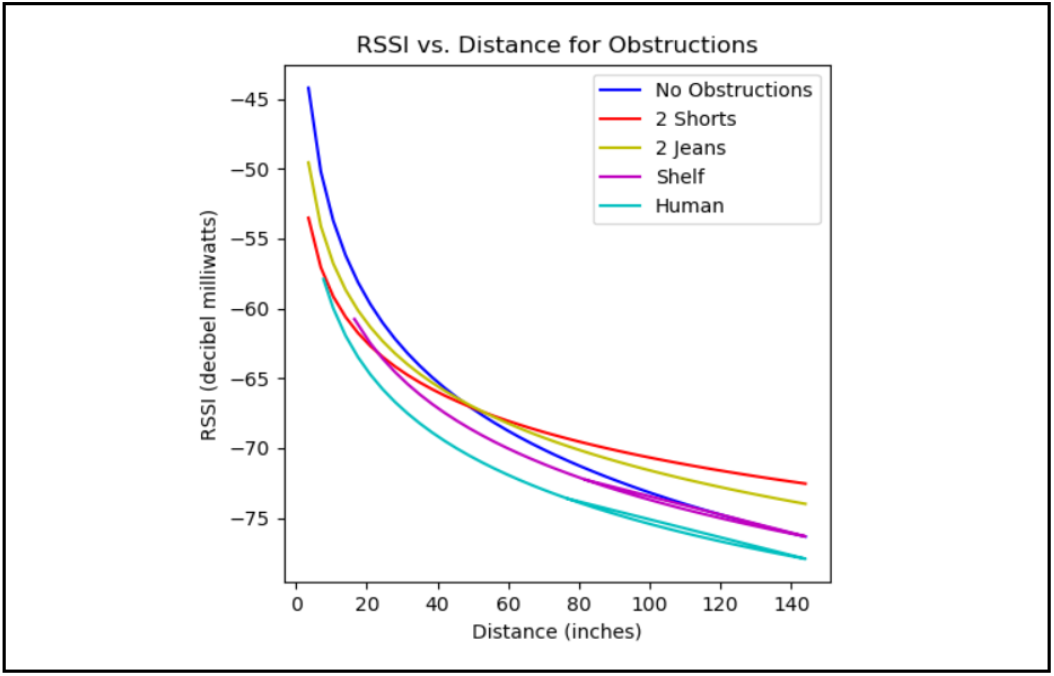
RSSI values for exp. 1, 4, 7, 8, and 9 against raw distance data.

**Fig. 4.**
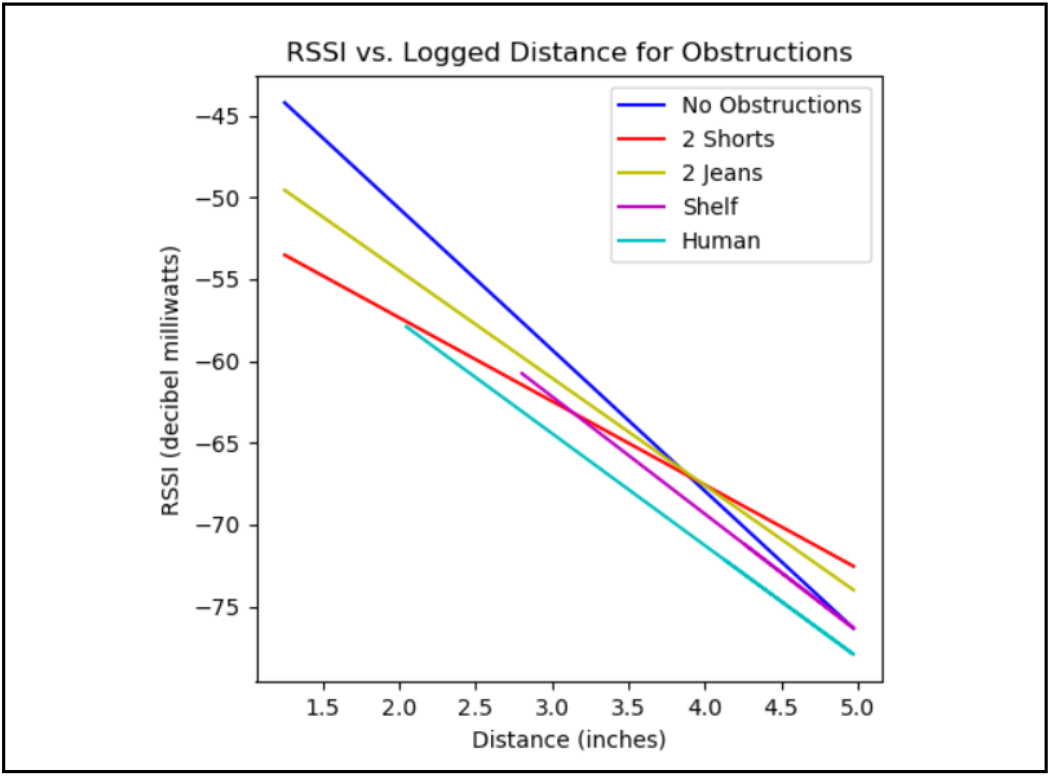
RSSI values for exp. 1, 4, 7, 8, and 9 against natural logged distance data.

It is apparent once again that a model from an experiment work with another experiment’s data. For example, exp. 7 being put through the models from exp. 1 and exp. 4 led to the exact same results. Surprising, exp. 9 with models from exp. 4 & 7 in table 3 and exp. 9 with models from exp. 7 & 8 in table 4 both have the same results, suggesting that perhaps temperature and humidity really do not have a large effect on predicting distance.

Given how I defined a success and better results, it seems that overall temperature and humidity not only did not help my prediction algorithm, but also made it worse and the number of false negatives has drastically increased. Looking at table 3, my algorithm successfully predicted true positives & negatives 66.813% of the time. If only a false negative count as a failure, then my model was successful 91.873% of the time.

## VI. Conclusions

### A. Hypothesis Evaluation

Hyp. 1 proved to be false. Only the physical obstruction of a human resulted in statistically significant results (table 2) when compared to exp. 1. Also, it is apparent from figures 3 & 4 that other obstructions do not make a significant difference. Therefore, common clothes like athletic shorts or jeans and a shelf in fact do not decrease the RSSI value enough to be considered significant. Moreover, the models of different experiments could be applied on another experiment, meaning that the data wasn’t that different from each other. For hyp. 2, although based on [3] atmospheric obstructions affect RSSI in a way to help better estimate distance, based on my results, hyp. 2 proved to be false. Due to the fact that the same result were created from experiment 9 under a variation of models, atmospheric obstructions most likely do not make a significant impact on the predictions made with my data. Also, according to table 4, more severe prediction errors were made when such obstructions were taken into account of. There were too many false negatives, suggesting that by adding temperature and humidity, my algorithm would overestimate the distance. Thus, atmospheric obstructions do not improve proximity prediction.

Given the success rates, hyp. 3 proved to be correct. Not noly was an algorithm constructed, a decent algorithm was successfully created from Raspberry Pi signals with a fair success rate. Although the algorithm is not perfect, it is acceptable. Also, hyp. 4 proved to be true as table 2 shows statistically insignificant results between experiment 2 & 3 and 5 & 6. Therefore, it doesn’t matter which Raspberry Pi is experiencing the obstruction.

### B. Noteworthy Conclusions

It seems that putting an obstruction on the scanner pi resulted in more of a decrease in RSSI value than the advertiser pi. From table 2, the test of whether the scanner pi had a larger effect or not was proven to be statistically significant for the obstruction of jeans (exp. 5 & 6). Although it was not statically significant for shorts, the p value for the test of if the scanner pi has more effect was more significant than the test of if the advertiser pi has more effect (0.447 < 0.553). Additionally, for shorts and jeans, having one obstruction on only one pi was not statistically significant compared to having one obstruction on both pis, as shown on table 2. Adding on, due to the nature of the RSSI and distance’s relationship, it there is a sudden jump in data, it is likely to concluded that an extremely large and dense object has been placed in between two devices, or the two devices have suddenly approached each other and entered a dangerously close distance. Another noteworthy conclusion is that RSSI most likely cannot be reliably be used to determine proximity as there are many other factors in a specific environment needed to be taken into account. Even though I experimented in a very controlled environment, I was not able to get excellent results via my algorithms, so Bluetooth based contact tracing, may not be the most accurate.

### C. General Lessons Learned

Based on my results, Bluetooth-based contact tracing is not reliable in the real world as there are too many errors in predictions. Moreover, it would not be too feasible for everyone to be carrying a Raspberry Pi with them wherever they go as I did my research on the Bluetooth signals transmitted from one Raspberry Pi to another. Finally, I am hesitant to believe that the Bluetooth signals from Raspberry Pis are much different from the Bluetooth signals from smartphones. Therefore, analyzing Bluetooth signals from Raspberry Pis is not a better solution, as the accuracy is not superb enough to outweigh the other downfalls.

## VII. Next Steps

First off, I will repeat all experiments to confirm the accuracy of my current values and to better predict proximity. Second of all, I will measure just the effects temperature and humidity individually without the effect of distance and build a model from there to better test hyp. 2. I never solely measured those two, so I may get promising results afterwards. Thirdly, I’ll measure the effect of more obstructions as people do not only wear athletic shorts and jeans year-round. Specifically, I’d measure the effects of khaki shorts and long pants, winter coats, hoodies, etc... Fourthly, I would like to add on confidence intervals into my models to allow a better range for my predictions, hopefully resulting in better results. Finally, to bring this into the outside world, I will determine a method to implement Bluetooth signals from Raspberry Pis into a portable item for feasible use.

